# Multiple hypothesis correction is vital and undermines reported mtDNA links to diseases including AIDS, cancer, and Huntingdon’s

**DOI:** 10.1101/015602

**Authors:** Iain G. Johnston

## Abstract

The ability to sequence mitochondrial genomes quickly and cheaply has led to an explosion in available mtDNA data. As a result, an expanding literature is exploring links between mtDNA features and susceptibility to, or prevalence of, a range of diseases. Unfortunately, this great technological power has not always been accompanied by great statistical responsibility. I will focus on one aspect of statistical analysis, multiple hypothesis correction, that is absolutely required, yet often absolutely ignored, for responsible interpretation of this literature. Many existing studies perform comparisons between incidences of a large number (*N*) of different mtDNA features and a given disease, reporting all those yielding *p*-values under 0.05 as significant links. But when many comparisons are performed, it is highly likely that several *p*-values under 0.05 will emerge, by chance, in the absence of any underlying link. A suitable correction (for example, Bonferroni correction, requiring *p* < 0.05/*N*) must therefore be employed to avoid reporting false positive results. The absence of such corrections means that there is good reason to believe that many links reported between mtDNA features and various diseases are false; a state of affairs that is profoundly negative both for fundamental biology and for public health. I will show that statistics matching those claimed to illustrate significant links can arise, with a high probability, when no such link exists, and that these claims should thus be discarded until results of suitable statistical power are provided. I also discuss some strategies for responsible analysis and interpretation of this literature.

## Introduction

In the interests of making this communication suitable for a general audience, I hope that expert readers will forgive a basic introduction. Mitochondrial DNA (mtDNA) is a molecule that encodes important aspects of the cellular machinery required for mitochondrial functionality in eukaryotic cells. MtDNA encodes this machinery through a sequence of nucleotides, chemical units that are often represented by their initial letters *A*, *C*, *G* and *T*. This nucleotide sequence is interpreted by the cell as a set of instructions for producing proteins and other components of the mitochondrion. MtDNA is subject to mutations, giving rise to variability in mtDNA sequences in individuals and across populations. Mutations in mtDNA can involve a replacement of one nucleotide by another (for example, an *A* becoming a *G*), insertions or deletions of sets of nucleotides (for example, an *A* being omitted from a sequence), and others which are of less concern in this communication.

Mitochondria are central sources of cellular energy, and dysfunction in mitochondria is linked to many diseases [1]. Mutations in mtDNA can result in the incorrect production of mitochondrial machinery and thus cause human disease. An example is the *A*3243*G* mutation (also written in forms including 3243*A* > *G*, *mt*3243*A* > *G*, 3243*A*/*G*, and others; to be read as ‘a change at position 3243 in mtDNA from *A* to *G*), which often causes the inherited disease MELAS [2]. Many other disease-linked mtDNA mutations have been reported; the ability to sequence mtDNA cheaply and quickly has led to a common recent research theme seeking links between such mutations and disease.

A recent, large-scale analysis of biomedical literature found that most published research is wrong [3]. It is important to realise that this statement is not provocative hyperbole, but is a quantitative claim, substantiated by a large scale metaanalysis, showing that statistical errors and misdemeanours mean that over 50% of reported results are incorrect. This profoundly disturbing finding is anecdotally supported by reports of the lack of repeatability in papers considered ‘landmarks’ in cancer science ([4]; only 6 of 53 papers could be reproduced) and more general drug design ([5]; only about 25% of published preclinical results could be appropriately validated). Many statistical and scientific issues contribute to this state of affairs; in this communication I focus on one statistical problem, multiple hypothesis correction (reviewed in, for example, Ref. [6]), in one particular field, identifying mtDNA links to disease. I will demonstrate how seemingly significant results can arise by chance when multiple hypotheses are not corrected for, briefly show how these corrections can be applied, and discuss a (non-exhaustive) set of recent results which should be discarded due to their absence of such correction. I conclude by urging authors and reviewers to avoid the incorrect and unethical neglect of multiple hypothesis correction, and readers to bear this vital approach in mind.

## Results

### Incorrect results arise from association studies without multiple hypothesis correction

A typical study in this field will examine a set of patients and a set of controls, and seek links between specific mtDNA features and disease in these groups. For example, a recent paper [7] has reported that a mutation at site 16290 in human mtDNA is significantly linked to AIDS prevalence. The line of reasoning is as follows. 18 people possess this mutation: of these, 9 have AIDS and 9 do not. 220 people do not possess the mutation: of these, 61 have AIDS and 159 do not. Based on these figures, the odds of having AIDS given the mutation (9/9) are higher than those of having AIDS without the mutation (61/159). However, it is not unreasonable to think that these figures could have arisen purely through chance, with no connection between the mutation and AIDS prevalence. To provide support for the existence of a real connection, we need a way to quantify how likely such figures are to arise under this ‘null hypothesis’ that no connection exists.

The way this likelihood is often presented is based around the concept of a ‘*p*-value’. A *p*-value generally represents the probability that an observation at least as extreme as the one observed could arise under a null hypothesis. In the context of these association studies, a *p*-value gives the probability with which we expect to see a difference in odds as high as the one we actually observe, if no link exists between mtDNA feature and disease. Thus, a *p*-value of 0.05 corresponds to a 5% probability that the result we have observed could have arisen by chance without any underlying link. This threshold of 5% is often deemed ‘significant’ *for a single observation.* Statistical analyses including Pearson’s chi-squared test and Fisher’s exact test [8] are typically employed to estimate the *p*-value associated with seeing the observed figure under the null hypothesis of no link. In the case above (9/9 and 61/159), Pearson’s chi-squared test gives a *p*-value of *p* ≃ 0.046, which the authors report as the signature of a significant link.

However, we must reflect on what a *p*-value means when we make many observations. Recall that the *p*-value reflects the probability with which an observation can arise when no link exists; *we must thus expect to see p-values of 0.05 or under around* 5% *of the time even when no link exists.* For example, if we investigate 1000 features, none of which are linked to the disease of interest, we would still expect to see around 50 features with *p* < 0.05 arising purely by chance. An individual observation of *p* < 0.05 is thus (much) more common when multiple comparisons are performed, and such an observation provides little or no evidence against the null hypothesis (captured, for example, in Ref. [9]).

This brings us to the key message of this communication: *The p* < 0.05 *‘significance’ condition cannot be applied over more than one comparison without correcting for the number of comparisons involved.* Because of this, many reported links between disease prevalence and individual SNPs or other mtDNA features cannot be regarded with the ‘significance’ that the authors claim.

To demonstrate this problem, I present a simulation study. *S* = 1000 synthetic datapoints are constructed, each corresponding to a ‘patient’. *N* = 25 mtDNA features will be considered in each patient. Each of these *N* features is randomly chosen, either mutated (with probability *µ* = 0.2) or wildtype (with probability 1 – *µ*). Each patient is then randomly categorised as diseased (with probability *σ* = 0.2) or healthy (with probability 1 – *σ*), completely independently of any genetic feature. I then use Pearson’s chi-squared approach, as in Ref. [7], to seek links between genetic features and disease, with the knowledge that no such links in fact exist. A typical set of results is present in Table 1, where we observe two *p*-values under 0.05. In Fig. 1A we see that it is very common for one or more *p*-values under 0.05 to appear even when there is no link between any genetic feature and the disease in question. To summarise the problem: an uncorrected analysis of multiple comparisons is likely to yield (many) false positive results.

**Table 1:**
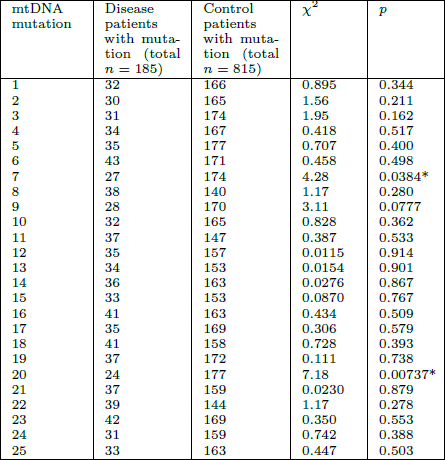
False positive results from applying a *p* < 0.05 significance criterion over multiple comparisons without correction. Results from a synthetic study investigating the link between different mtDNA mutations and disease prevalence (see text). Mutations and disease are randomly assigned and are in no way linked; however, due to the large number of comparisons, we observe some associated *p*-values under 0.05 (asterisks; associated with mutations 7 and 20). These do not signal any link between mutation and disease (none exists) but arise due to chance. Multiple hypothesis correction must be employed to avoid erroneously labelling these links as ‘significant’. This is not an unlikely, cherry-picked example; Fig. 1A shows how often we can expect such observations due to chance.

**Figure 1:**
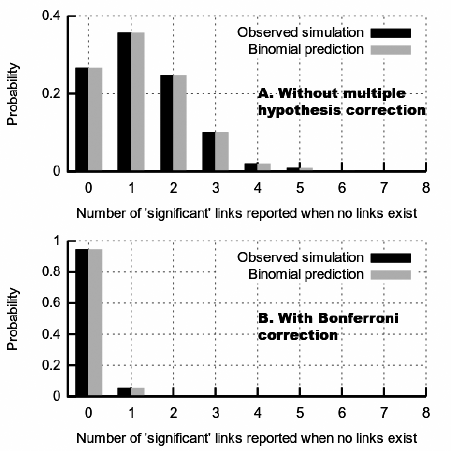
Several *p*-values under 0.05 should be expected when performing multiple comparisons, even when no significant link exists. **A.** The probability of observing a given number of *p*-values under 0.05 when performing 25 comparisons, none of which is associated with a real link. Results are presented both for the theoretical expectation and for the simulation experiment in Table 1 and described in the text. There is a 72% chance that we observe at least one ‘significant’ result despite the fact that none exist; clearly, we need to correct for this effect. **B.** Bonferroni correction in this case requires *p* < 0.05/25 for a result to be labelled as ‘significant’. The probability of reporting a false significant link having corrected for multiple hypotheses is now a more reasonable 0.05.

For example, in Ref. [7], the authors consider 25 mtDNA mutations. They thus explore 25 possible links between genetic features and AIDS, and find *p*-values under 0.05 for two of these features. These are then interpreted as significant evidence against the null hypothesis. But as 25 different experiments have been performed, we should expect to see *p*-values under 0.05 arising just by chance (as in Table 1). In fact, there is a 72% chance in this case that at least one *p*-value under 0.05 will arise, and a 23% chance that will we see exactly two, as the authors do, under the null hypothesis (Fig. 1A). The appropriate probabilities – that of observing a given number of *p*-values under 0.05 – arise from a binomial distribution with *p* = 0.05 and *N* equal to the number of comparisons (25 in this case), which can straightforwardly be visualised and explored using software like the Caladis probabilistic calculator [10].

The reader will note that the synthetic case in Table 1 displays comparable (in fact, stronger) evidence than Ref. [7] for the existence of specific links, despite the fact that no such links exist. This problem arises because multiple comparisons have not been suitably accounted for, and without further evidence, the claimed links between mtDNA and AIDS of Ref. [7] must be regarded as unsupported.

It is important to note that these results are a general property of statistical tests interpreted through *p*-values, and are not a consequence of a particular choice of methodology (for example, Pearson’s chi-squared test). Multiple comparison *p*-values from, for example, linear regressions also need to be corrected. In the simulation example, the results do not depend on choice of parameterisation; different *µ*, *σ,* and *S* will give comparable results, and I have deliberately chosen reasonably large sample sizes (*S* = 1000) to illustrate that the necessity for multiple hypothesis correction is not removed through increased sample size. To summarise, multiple hypothesis correction is not an optional statistical nicety that can be employed if and when it is desired; it is of absolute importance to avoid the reporting of spurious false positive results.

### Multiple hypothesis correction is simple and should be ubiquitous

Fortunately, methods for correcting this multiple comparison problem exist, and there is a substantial literature on the subject (Ref. [6] gives a comprehensive review). Broadly, these methods involve an adjustment of the definition of ‘significance’ to reflect the number of comparisons that have been performed. To link with the existing statistical literature and textbooks, a slightly more formal nomenclature must be adopted. A false rejection of the null hypothesis, as demonstrated above (a false positive result) is a Type I error; a false negative result is a Type II error. The probability of at least one Type I error is known as the family-wise error rate. The expected proportion of Type I errors is known as the false discovery rate. The probability that the proportion of Type I errors exceeds a certain value is known as the false discovery exceedance.

Correction methods seek to control the family-wise error rate, the false discovery rate, or the false discovery exceedance. Bonferroni correction is perhaps the best known of these approaches. A Bonferroni correction using the 0.05 significance level involves regarding *p*-values as signals of a significant departure from the null hypothesis only when *p* < 0.05/*n*, where *n* is the number of comparisons performed. This aims to ensure that the family-wise error rate does not exceed 0.05. The application of Bonferroni correction to the earlier simulation example confirms that this criterion is satisfied (Fig. 1B), demonstrating the dramatic reduction of false positive reports compared to the case where multiple hypothesis correction is absent.

Bonferroni correction is viewed as quite conservative, and its strict nature may lead to Type II errors. Arguably, statistical conservatism is no bad thing in a research climate where more than half of published results are incorrect [3], but alternative correction strategies exist to reduce the probability of Type II errors, and can be employed as long as they are compatible with the structure of the scientific study. The names of some methods controlling the family-wise error rate include Bonferroni, Holm, Hochberg, and Sid´ak; procedures controlling the false discovery rate include Benjamini-Hochberg or are often simply referred to by the acronym FDR. The purpose of this communication is not to describe and review these methods (a task performed, for example, in Ref. [6]), but to urge the reader to seek evidence of multiple hypothesis correction in interpreting mtDNA studies.

### Many recent studies present unsupported results linking mtDNA with disease

We have seen that the results from Ref. [7], reporting links between mtDNA mutations and AIDS prevalence, can easily result by chance under a null hypothesis of no links, and must therefore be discarded until stronger confirmatory evidence is provided. This study is by no means unique in the literature: here I examine a small set of other studies in which the statistical methodology must be questioned. These examples are drawn simply from examining a set of search results for mtDNA associations with disease from recent publications and is certainly not exhaustive.

Ref. [11] analyses 35 SNPs seeking links with Huntingdon’s disease. The authors report 8 SNPs that display *p* < 0.05. An additional problem exists with this study in that authors do not quote exact *p*-values, rather just noting those values below 0.05. It is therefore impossible to immediately interpret whether the authors have found 8 *p*-values of 0.049, all of which Bonferroni correction would discard, or 8 *p*-values of 10^−16^, all of which would remain ‘significant’ under Bonferroni. A re-analysis of their data using Fisher’s exact test shows that all but two of the reported SNPs should be discarded under Bonferroni correction (see below).

Two other examples focus on mutations in the D-loop region of mtDNA linked to ovarian cancer [12] and non-Hodgkin lymphoma [13]. Disturbingly, these studies seem to be representative of a set of similar studies, in a variety of journals, on links between mtDNA D-loop features and various cancer incidences, all of which fail to employ multiple hypothesis correction [14, 15, 16]. These studies follow a very similar core methodology, identifying a set of SNPs in a patient and control cohort, focussing on a set of SNPs where the rare allele is present in more than 5% of controls or patients, and seeking links between this set and the cancer of interest. Ref. [13] gives the size of the set of SNPs considered as 26; Ref. [12] apparently omits this important information. Both studies then report SNPs with *p*-values under 0.05 as significantly linked to their respective cancer types without multiple hypothesis correction. Furthermore, both studies claim a *p*-value of zero for some SNPs, making the claim that there is zero probability that their observed links could have arisen by chance. This is an impossible statement under any reasonably constructed null hypothesis (although *p*-values may be extremely small) and makes it impossible for the reader to apply the required multiple hypothesis correction themselves.

Although absence of important data in some of the above papers prevents a full re-analysis, some consequences of Bonferroni correction are immediately clear. Such a re-analysis can straightforwardly be applied by the reader by multiplying each *p*-value (stated without multiple hypothesis correction) by the number of mtDNA features examined, and checking if the result remains under 0.05. For example, correcting a *p*-value of 0.01 to account for a study of 20 SNPs would give 0.20, which is over 0.05 and thus discarded. Following Bonferroni correction, neither mtDNA haplogroup A, nor specific mutations at sites 16209 or 16319, can be linked to AIDS [7]. *A*263*G* cannot be linked to non-Hodgkin lymphoma; *G*200*A*, *C*16362*T*, *A*249*del* cannot be linked to diffuse large B-cell lymphoma; *C*315*insC* cannot be linked to T-cell lymphoma [13]. None of *G*207*A*, *C*523*del*, *T*254*G*, *A*259*G*, *C*418*G*, *C*441*A*, *C*476*A*, *C*530*T*, *A*249*del*, *A*263*G* can be linked to ovarian cancer [12] (there is also an issue in this study with the use of Pearson’s chi-squared test with low observation counts). None of *C*16069*T*, *T*16126*C*, *T*16189*C*, *C*16223*T*, *T*16086*C*, *C*16150*T* can be linked to Huntingdon’s disease [11]. Multiple hypothesis correction also removes support for SNPs described as significantly connected to gastric cancer [16], breast cancer [15]; other erroneous reports no doubt exist in the broader literature.

These and similar corrections do not mean that no link exists, but rather that statistical support for such a link is not yet evident. In some of the above studies, a subset of reported links between some proposed mtDNA features and disease do survive multiple hypothesis correction and can therefore be subjected to scientific scrutiny with less concern that they represent statistical artefacts. Large-scale and rigorous analyses (such as a recent examination of mtDNA links to cancer [17]) is desirable to test these hypotheses appropriately.

## Discussion

This communication has focused on a particular aspect of statistical methodology frequently employed in mtDNA studies. A discussion of this topic should naturally be framed in a broader discussion of the merits and shortcomings of analyses based upon *p*-values [18], a focus on statistical significance as opposed to the size and importance of effects [19], and frequentist and Bayesian approaches to multiple hypothesis testing [20]. Space in this communication limits such a discussion but the references above and references therein may provide valuable context.

Working in the paradigm of frequentist tests for mtDNA-disease association, we have seen that multiple hypothesis correction is vital to avoid reporting erroneous links between mtDNA and disease. A range of correction strategies exist and can be applied; but an appropriate form of multiple hypothesis correction *must* be performed in analysing the results of these studies. To fail to do so is not just scientifically incorrect, but is unethical and runs a tangible risk of misguiding biological and medical research. Readers should be aware that multiple hypothesis correction is necessary and seek evidence of one of the above (or other) procedures having been applied; and should discard reports resembling those in Table 1 which are uncorrected and cannot be regarded as demonstrating any link, even a ‘trend’ or ‘suggestion’, between mtDNA and disease.

### Specific guidelines

To avoid the reporting of false positive results and to promote reproducible analyses in association studies, the following information should be included when applying frequentist tests for the presence of relationships or links between factors.

(i) A description of all the comparisons performed, including the total number and the statistical approach employed.

(ii) The source data and a measure of relationship strength (for example, an odds ratio), ideally for each comparison, and certainly for each comparison claimed to be significant, such that the statistical approach can easily be reproduced and the magnitude of reported effects can be assessed.

(iii) Precise *p*-values, ideally for each comparison, and certainly for each comparison claimed to be significant. The reporting of inequalities alone (for example, *p* < 0.05) should be discouraged, as this prevents subsequent re-analysis of the results. If a *p*-value is so low that computational issues exist in obtaining a precise estimate, an upper bound inequality can be used (for example, *p* < 10^−16^).

(iv) A description of an appropriate multiple hypothesis correction protocol applied to the results (for example, Bonferroni correction). This important inclusion is the focus of this communication.

(v) A definition of significance given the correction in (iv) (for example, *p* < 0.05/*n* for Bonferroni correction with *n* comparisons), and a list of those results that fulfil this criterion.

